# REVEALS: An Open Source Multi Camera GUI For Rodent Behavior Acquisition

**DOI:** 10.1101/2023.08.22.554365

**Authors:** Rhushikesh A. Phadke, Austin M. Wetzel, Luke A. Fournier, Mingqi Sha, Nicole M. Padró-Luna, Alberto Cruz-Martín

## Abstract

Understanding the rich behavioral data generated by mice is essential for deciphering the function of the healthy and diseased brain. However, the current landscape lacks effective, affordable, and accessible methods for acquiring such data, especially when employing multiple cameras simultaneously. We have developed REVEALS (**R**odent B**E**ha**V**ior Multi-cam**E**r**A L**aboratory Acqui**S**ition), a graphical user interface (GUI) written in python for acquiring rodent behavioral data via commonly used USB3 cameras. REVEALS allows for user-friendly control of recording from one or multiple cameras simultaneously while streamlining the data acquisition process, enabling researchers to collect and analyze large datasets efficiently. We release this software package as a stand-alone, open-source framework for researchers to use and modify according to their needs. We describe the details of the GUI implementation, including the camera control software and the video recording functionality. We validate results demonstrating the GUI’s stability, reliability, and accuracy for capturing and analyzing rodent behavior using DeepLabCut pose estimation in both an object and social interaction assay. REVEALS can also be incorporated into other custom pipelines to analyze complex behavior, such as MoSeq. In summary, REVEALS provides an interface for collecting behavioral data from one or multiple perspectives that, combined with deep learning algorithms, will allow the scientific community to discover and characterize complex behavioral phenotypes to understand brain function better.

## INTRODUCTION

The field of neuroscience has witnessed significant technological advancements in the last two decades (Bargmann & Newsome, 2014; Bassett et al., 2020). Researchers now have access to sophisticated tools and equipment that allow for more precise and detailed measurement of neural activity and behavior in rodents (Aharoni et al., 2019; Ghosh et al., 2011; Huang et al., 2021; Markowitz et al., 2018; Marks et al., 2022; Mathis et al., 2018, 2020; Wiltschko et al., 2015). As our understanding of the brain and its complex neural circuits deepens, researchers are interested in investigating the relationship between specific neural circuits and behavior. This necessitates the development of more complex behavioral experiments – and robust means to consistently capture these experiments for subsequent analysis – to target and manipulate brain regions or pathways to study their specific role in behavior (Bargmann & Newsome, 2014; Bassett et al., 2020). However, the field currently lacks a unified, easy-to-use, turn-key GUI-based application that will permit researchers to capture behavioral videos dependably and necessitates only little prior technical knowledge. Instead, this forces research groups to independently develop their own methods of video acquisition or rely on programs such as Bonsai (https://bonsai-rx.org), Micro-manager (https://micro-manager.org), and Spinnaker (https://www.flir.com), each of which harbors inherent issues.

Researchers are also increasingly interested in studying complex rodent behaviors relevant to human psychiatric or neurological disorders (Huang et al., 2021; Johnson et al., 2022; Sriram et al., 2020), which requires the design of behavioral experiments that can capture and assess multiple behavioral dimensions simultaneously in an unbiased manner (Johnson et al., 2022; Markowitz et al., 2018; Marks et al., 2022; Mathis et al., 2020). Importantly, these data can potentially be translated into biomarkers of disease (Ewen et al., 2021; Hidalgo-Mazzei et al., 2018; Jacobson et al., 2019).

Complex behavioral experiments have the potential to allow researchers to model and investigate human behaviors or cognitive processes more accurately. Multiple cameras placed strategically in the experimental setup can help achieve this goal (Del Rosario Hernández et al., 2023), allowing for comprehensive observation of the rodents from different angles and perspectives. This multiple-perspective setup enables researchers to capture various behavioral parameters and provides a more complete understanding of the animals’ actions, permitting more detailed analysis. Separately, multiple cameras can also provide redundancy in data collection; if one camera fails or misses a critical event, other cameras can compensate to ensure that the data collection is not compromised. This redundancy minimizes the risk of losing valuable information due to technical problems or human error.

However, researchers today lack effective, affordable, and accessible approaches for obtaining such behavioral data, particularly when utilizing multiple cameras simultaneously. To address this issue, we have created REVEALS (**R**odent B**E**ha**V**ior Multi-cam**E**r**A L**aboratory Acqui**S**ition), a graphical user interface (GUI) for acquiring rodent behavioral data via one or multiple USB3 FLIR cameras. REVEALS facilitates user-friendly management of one or several concurrent recordings from numerous cameras, streamlining the data collection process and empowering researchers to efficiently amass and analyze extensive datasets of rodent behavior.

We made this software package available as an independent, open-source framework that researchers can freely utilize and adapt to suit their specific research requirements. In this article, we elaborate on the technical aspects of the GUI implementation, encompassing camera control software, video recording capabilities, and synchronization mechanisms used to integrate distinct camera feeds. We substantiate the GUI’s robustness, dependability, and precision in capturing and analyzing rodent behavior through validation experiments employing DeepLabCut (DLC) pose estimation in object and social interaction experiments. REVEALS can be seamlessly integrated into existing DLC (Mathis et al., 2018) and MoSeq (Markowitz et al., 2018; Wiltschko et al., 2015) workflows to analyze intricate rodent behavior.

In summary, REVEALS offers an intuitive interface for gathering behavioral data from diverse viewpoints. When coupled with deep learning algorithms, it is a powerful tool for discovering and characterizing previously unidentified behavioral traits, thereby enhancing our comprehension of both healthy and diseased brain functionality.

## METHODS

### Ethics statement

All experimental protocols were conducted according to the National Institutes of Health (NIH) guidelines for animal research and were approved by the Boston University Institutional Animal Care and Use Committee (IACUC; protocol #17-031).

### Animals

All mice were group housed on a 12-hr light and dark cycle with the lights on at 7 AM and off at 7 PM and with food and water *ad libitum*. Mice used in the object and social interaction assays included 6 to 12 weeks old C57BL/6J (Jackson Laboratory, strain #: 000664, RRID:IMSR_JAX:000664) and CD-1 (Charles River Laboratories, strain code: 022, RRID:IMSR_CRL:022) mice.

### Behavioral Assays

Object and social interaction assays were run in an otherwise-empty arena (46 cm x 23 cm). In the object interaction assay, an object made from two 6-well plates was temporarily secured with a magnet to a specific end of the arena (randomly selected) and mice were free to explore for 2 min. In the juvenile interaction assay, a C57 and a juvenile CD1 mouse were placed in the arena together and were free to interact for 2 min.

### Behavioral analysis

All behavior was analyzed using DLC (Mathis et al., 2018), an open-source software package that uses deep neural networks to automatically track body parts from videos, as described in Comer et al. (Comer et al., 2020) and Johnson et al. (Johnson et al., 2022). We confirmed accurate tracking of mice based on this software by close inspection of videos once they had been annotated by DLC.

## RESULTS

The usage of the REVEALS GUI and associated pipeline proceeds in 5 to 6 steps (**Fig. 1**). The first part of the application controls the camera setup and recording of videos using either one or multiple cameras. Each recording session generates a series of videos named “*behavcam_[video number].avi”* and a timestamp file to track the writing time of every frame in these videos. These videos can then be run through the DLC or MoSeq pipelines, or a combination, resulting in frame-by-frame tracking of body part location, as demonstrated later in this manuscript. Combining these outputs with the timestamp files will allow the user to account for any frame mismatches that may occur when recording from multiple FLIR cameras, thus allowing for complete synchronization of tracking data which can be further analyzed using custom codes to extract behavioral paradigms.

**Figure 1.**
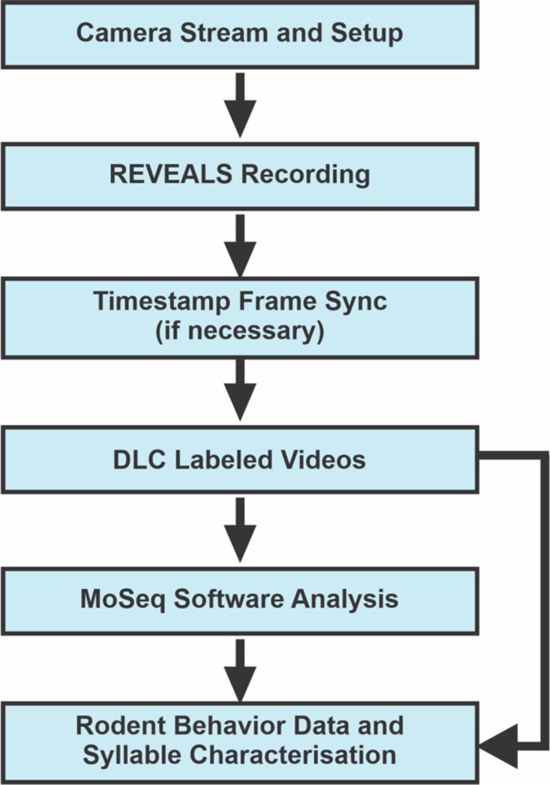
Overall workflow of rodent behavior analysis beginning with REVEALS multi-camera video capture. However, REVEALS is compatible with other custom-made analysis pipelines.

### Opening the REVEALS GUI

REVEALS is a stand-alone software solution that allows the user to obtain either single or time-synched multi-camera recordings of behavioral data (Fig. 1). To work with REVEALS, the user must first download and install Anaconda 3 (*Anaconda Software Distribution. (2020). Anaconda Documentation. Anaconda Inc.*, 2020). The user can find detailed step-by-step installation instructions in a Readme.md file on our GitHub page (https://github.com/CruzMartinLab/REVEALS). There are two ways to install REVEALS depending on if the user wants to use the base packaged version or make custom changes to the script to better suit their specific research questions. The details on installing the packaged version can be found on the linked GitHub page; separately, below we describe the usage of the python script version.

Anaconda3 is an open-source, cross-platform, language-agnostic package manager and environment management system. After installation, to use REVEALS, the user must open the Anaconda command prompt window and call the Conda environment using the command:

`conda activate reveals`

This command allows all required packages and installations to become active. After this command, the user will find the directory where the python script for the REVEALS pipeline is saved using the command:

`cd [PATH WHERE INSTALLATION WAS MADE]`

The user can then open the REVEALS GUI as a final step using the command:

`python reveals.py`

### Camera Initialization

Opening the REVEALS GUI brings the user by default to the first of two tabs, *“Camera Setup”*. It is here where up to three cameras (tested, capacity for more than three cameras may depend on the specifications of the computer) can be recognized by REVEALS by pressing the “*Connect”* button (Fig. 2A). After pressing *“Connect”*, the number of cameras connected to the computer appears in the line *“Number of Cameras=”* at the top right of the *“Camera Setup”* tab. The cameras are arranged by serial number and organized numerically such that the lowest serial number corresponds to *“Camera 1”*.

**Figure 2.**
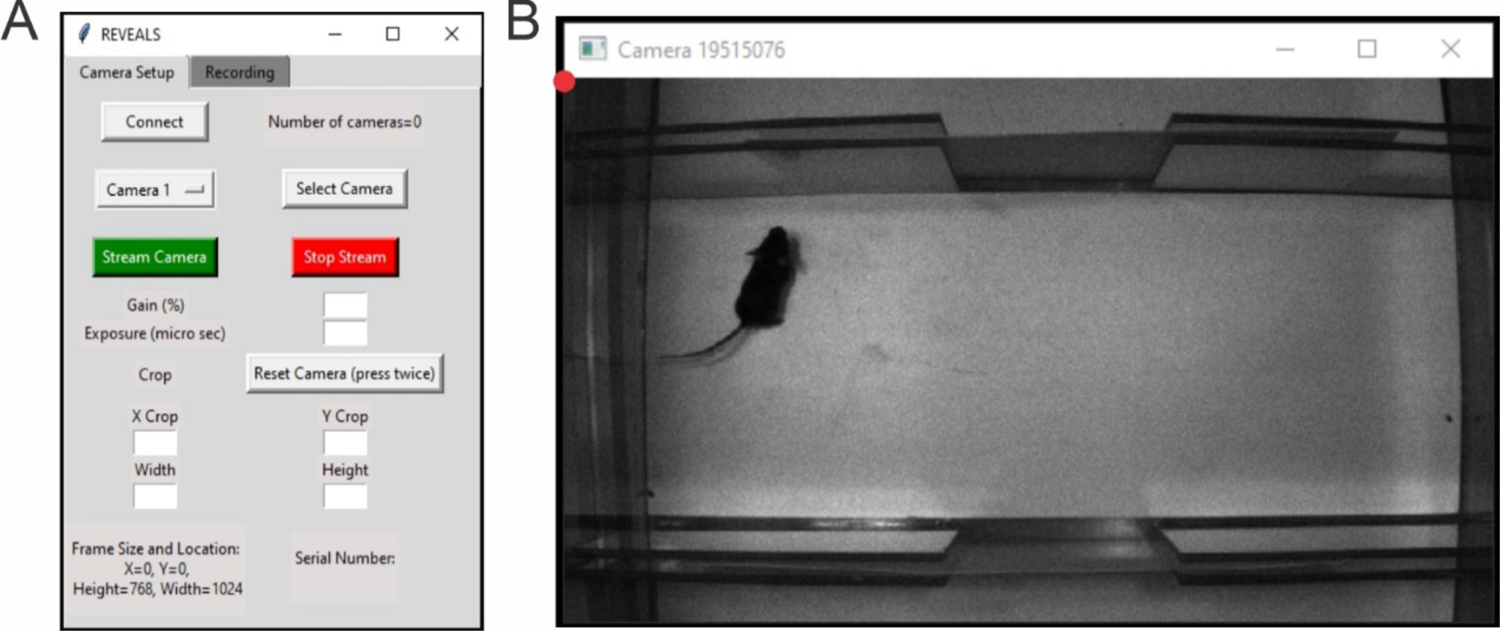
REVEALS camera setup and single camera streaming for the field of view. (**A**) The default window when the REVEALS application is opened. The GUI displays a *“Camera Setup”* and *“Recording”* tab. On the *“Camera Setup”* tab, the *“Connect”* button is used to discover the connected cameras. The *“Camera 1”* dropdown menu, *“Select Camera”*, *“Stream Camera”* and *“Stop Stream”* buttons are used for selecting a particular camera and streaming its view to ensure a stable connection as well as desired field of view. *“Gain (%)”* and *“Exposure (μs)”* input boxes are used to modify the camera’s internal light settings. *“X crop”*, *“Y crop”, “Height”* and *“Width”* input boxes are used to determine field of view. *“Reset Camera”* will reset the crop, gain, and exposure values to camera defaults. *“Serial Number”* label will display the serial number of the selected camera, while the *“Frame Size and Location*” label will keep track of the current field of view’s dimensions. (**B**) The live camera feed window when *“Stream Camera”* is pressed. The red dot represents the (0,0) point of the grid for cropping. The window name corresponds to the serial number of the selected/actively streaming camera.

The next step is to initialize each camera using the dropdown menu. When a given camera is chosen via the dropdown menu, pressing the *“Select Camera”* button will initialize that camera. By default, the dropdown begins with *“Camera 1”.* Upon pressing the *“Select Camera”* button, the fields to set crop values, gain percentage, and exposure (in μs) are displayed, along with the serial number of the selected camera.

### Selecting and Saving Camera Parameters for Recording

After selecting the desired camera, pressing the *“Stream”* button will bring forth a live feed from the selected camera. The gain, exposure, and crop values can be changed by entering the desired values into the respective text boxes in the *“Camera Setup” tab*. The user should only input positive crop values. Cropping in REVEALS utilizes a grid system whereby the top left corner of the camera feed represents the origin (0, 0) (Fig. 2B). By changing the *“X Crop*” value, the left side of the current camera feed will be cropped by the indicated amount, and the new cropped camera feed will span the length indicated by the *“Width”* value (also editable). Similarly, when the *“Y Crop”* value is changed, the top of the current camera feed will be cropped by the amount indicated, and the new cropped camera feed will span the distance of the *“Height”* value (also editable). It is easiest to adjust these crop values, as well as the gain and exposure, while having the camera actively streaming. However, if doing so, the display of the camera feed will not update unless values are within maximum specifications. The user should not change the gain value for a given camera to more than 100%. The user should consider that high gain will lead to noise in the system, while too high exposure time could increase the number of dropped frames.

Finally, to save the desired crop, gain, and exposure values set, the user can press *“Stop Stream”*. Pressing this button will also close the live camera feed. When the stream is stopped, that camera is fully initialized and ready for use. If the user wants to change any of these values, *“Stream Camera”* and *“Stop Stream”* will have to be clicked again to let the user set and save, respectively, the changed values.

### Recording with Multiple Cameras

If the user is using multiple cameras, after selecting and saving the recording parameters for a given camera, they must utilize the dropdown menu to choose the next camera and follow the same process as given in the previous section of this document to select and save their desired parameters. In this way, each camera can have entirely unique crop, gain, and exposure values. The *“Frame Size and Locations”* text in the bottom left corner of the *“Camera Setup”* tab shows the last streamed crop values for the selected camera. Note that in some cases, FLIR cameras might default to an intermediate value of crop settings, or the user might want to return all camera settings to default. In these cases, selecting the corresponding camera from the drop-down menu and pressing *“Reset Camera”* twice will cause that camera’s values to be set to default internal values.

### Recording

Once the user initializes all cameras to their desired parameters, they can click on the *“Recording”* tab of REVEALS. The *“Folder Name”* text box is used to name the folder within which the output of the recording (camera’s videos (.AVI) and timestamp (.CSV) files) will be saved (Fig. 3A). If there is no input to the *“Folder Name”* field, the name of this folder will default to the date and time of the folder’s creation (YYYY_MM_DD HH_MM). Next, the user can choose the desired acquisition sampling rate in frames per second (FPS) from the dropdown menu of either 15, 30, 45, or 60. REVEALS sets the default acquisition sampling rate at 30 FPS. Once the user sets the acquisition sampling rate, they must define the recording time in seconds under *“Recording Time (s)”*. After this, the user can click the *“Stream”* button, which will cause windows for each camera to appear, displaying the camera feeds with the recording parameters that the user set in the *“Camera Setup”* tab (Fig. 3B). Once the cameras start streaming, the user can still alter the recording time. However, the acquisition sampling rate must be set before streaming. If the user needs to change the acquisition sampling rate, they must press *“Stop Stream”* and then proceed to change this value. The cameras must be streaming before the recording can be started. Once the recording begins, the stream will no longer be active, and what will be displayed is a still shot of the feed from the last moment before the recording started.

**Figure 3.**
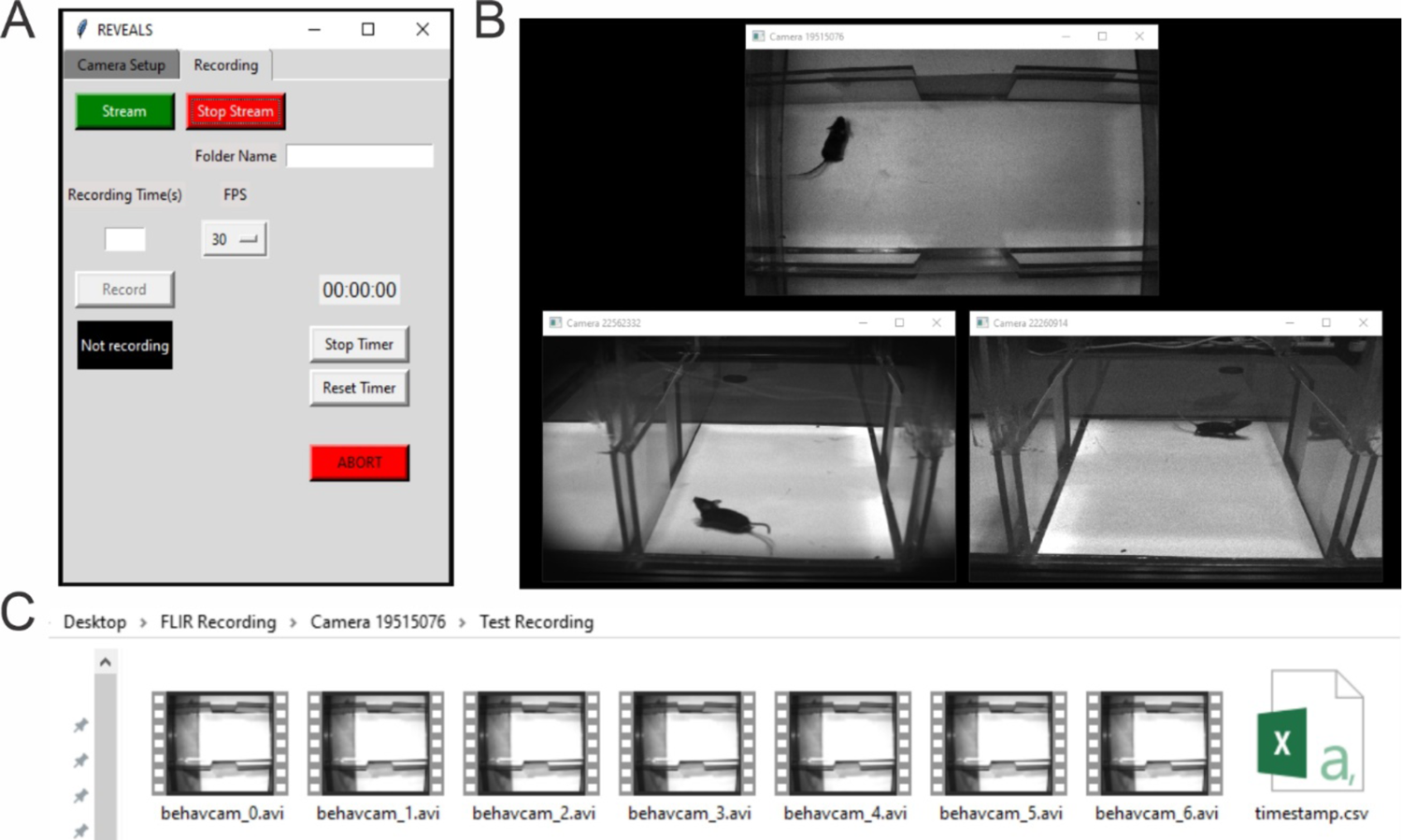
REVEALS recording setup and file organization. (**A**) The “*Recording*” tab of the GUI displays the “*Stream*” and “*Stop Stream*” buttons used to start/stop camera streaming before recording videos. “*Recording Time*”, “*FPS*”, and “*Folder Name*” options are available for the user to adjust recording parameters as desired. A timer on the right-side displays the time elapsed since the recording started, along with “*Stop Timer*” and “*Reset Timer*” buttons for further controlling this timer. The “*Record*” button is used to initialize the recording after streaming, with the label underneath showing the status of the recording. The “*ABORT*” button can be used to stop the recording at any point of time. (**B**) The live camera feed when *“Stream” is pressed, displaying live feeds of all connected cameras with the user-set camera parameters.* (**C**) An example folder for *“Camera 1”* after a 60 s recording at 60 FPS that holds the recordings and corresponding timestamp.csv file.

Once the user presses the *“Record”* button, a timer will indicate the current duration of the recording. Additionally, the text underneath the *“Record”* button will change from *“Not Recording”* to *“Recording”*. After the recording time has elapsed, this label will change to read *“Saving”* while all videos and timestamp files are saved. The recording process will be complete with all files saved once this label returns to *“Not Recording”*.

### Closing REVEALS

To close REVEALS, the user must close the streaming windows by pressing *“Stop Stream”*. Once the streams are stopped and closed, REVEALS can be closed by clicking the *“X”* in the upper right of the REVEALS GUI and selecting *“OK”* when prompted. If the user does not stop and close the streams before closing REVEALS, they can press the *“Cancel”* button when prompted to stop the streams.

### Finding Files

REVEALS will save the videos and timestamp files in the following path: “/Desktop/FLIR_Recordings/Camera [Camera Serial Number]” (Fig. 3C). Simply, within the folder titled, *“FLIR_Recordings”*, a folder for each connected camera is created. Inside of each of these folders belonging to each camera, the user will find sub-folders named either by the user (input by the user in the *“Recording Name”* field of the *“Recording”* tab or by the date and time of creation (default when there is no user input in the *“Recording Name”* field, see “Recording” section of this document above). By default, REVEALS creates videos containing 1000 frames; any remaining frames are saved as a shorter final video. For example, a recording with 4,500 frames will create five videos: four videos with 1000 frames, and the fifth with the remaining 500 frames. Within the timestamp .CSV file, REVEALS lists each frame with its corresponding time of occurrence in ‘ms’. The user can use the timestamp files later to synchronize the cameras’ videos if using multiple cameras.

### Performance Metrics for REVEALS

We performed multiple recordings at various sampling frequencies to determine the reliability of REVEALS (Fig. 4). The computer used to run the tests comprises 32 GB RAM and an Intel Core i7-7700 CPU with 3.60 GHz Quadcore processor. It has a 512GB NVMe SSD and 2TB HDD, with USB 3.0 ports. For a frame size of 750 by 1050 pixels, each recorded frame had a size of 230 KB. With these specifications, we found that REVEALS performed optimally at 15, 30 and 45 FPS, however performance declined while recording for more than 5 min at 60 FPS (**Fig 4A, B**). During a 5-min recording at 30 FPS (total of 9000 frames), REVEALS dropped only 0.12% of the frames. Thus, we recommend using 30 FPS for all whole body recordings that require a large field of view (Comer et al., 2020; Johnson et al., 2022) if the user is using a computer with less than 32 GB RAM (Fig. 4C, D). We have previously shown that recordings accomplished at 30 FPS are sufficient to capture the intricacies of whole body mouse behavior (Comer et al., 2020; Johnson et al., 2022). The performance loss at higher FPS can be remedied by increasing the memory, at which point REVEALS will perform equally well at 60 FPS.

**Figure 4.**
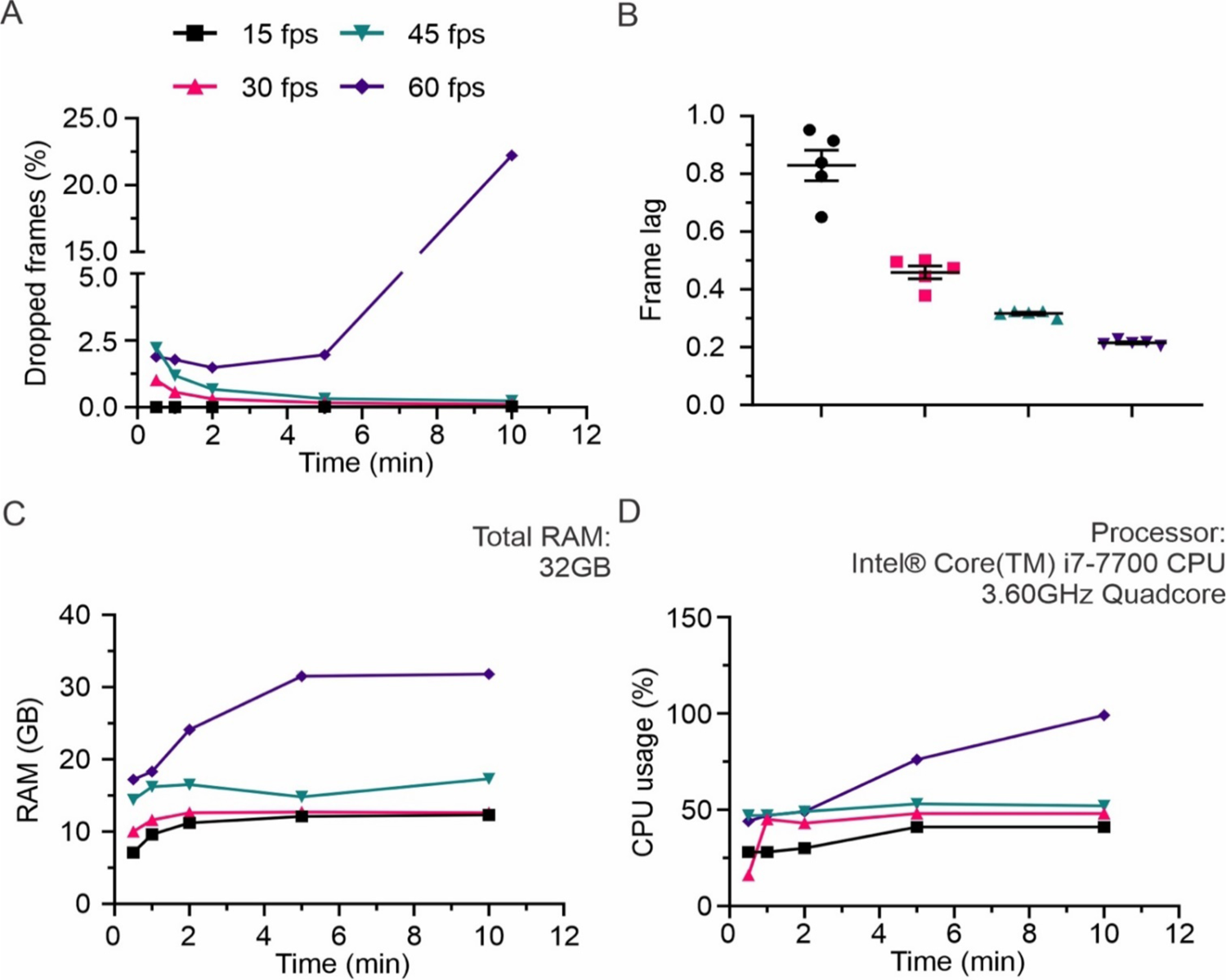
REVEALS performance and processor usage at 15, 30, 45 and 60 fps. (**A**) While recording at 15 and 30 FPS, REVEALS performed optimally, with an average of 0.2% of total frames dropped. At 45 and 60 FPS, there was an increased amount of frame drops, with an average of 1% and 5%, respectively. (**B**) The lag between recording timings of each frame for 3 cameras was approximately below 1 frame for all recording rates. N=5 recordings for each FPS. (**C**) For 15, 30 and 45 fps, the total memory usage throughout the duration of recording was approximately 14 GB. This showed an increase for 60 fps and was proportional to increasing recording time. (**D**) For 15, 30 and 45 FPS, the CPU usage throughout the duration of recording was approximately 42%. This showed an increase for 60 FPS and was proportional to increasing recording time. Graph in (**B**) show Mean ± SEM.

### Object and Social Interaction Tasks

The user can use REVEALS to obtain recordings with either a single camera or multiple cameras (**Supp.** Fig. 1, **Supp. Table 1**). Additionally, the user can feed these recordings to deep learning pose estimation pipelines such as DLC. To demonstrate this, we used REVEALS to perform multi-camera recordings during an object and social interaction task (Fig. 5). In the object interaction task (Fig. 5A**-D**), mice were free to explore an object for 2 min as their behavior was captured from three angles simultaneously. In an objection interaction task trial containing a total of 11 interactions (70 s of recording time), we showed that we could still reliably capture 100% of the interactions (red dots in Fig. 5D). However, in some of these instances, the snout or head of the mouse was occluded and not detected in one or two of the viewing angles, demonstrating the reliability and advantage of behavioral recordings using multicamera systems (Fig. 5D). Similarly, in the social interaction assay (Fig. 5E**-G**), a C57 and sex-matched CD-1 mouse were placed in the arena and their interactions were captured from three angles simultaneously. We employed DLC to consistently and precisely label the mice’s particular body parts, the object’s corners, and the arena.

**Figure 5.**
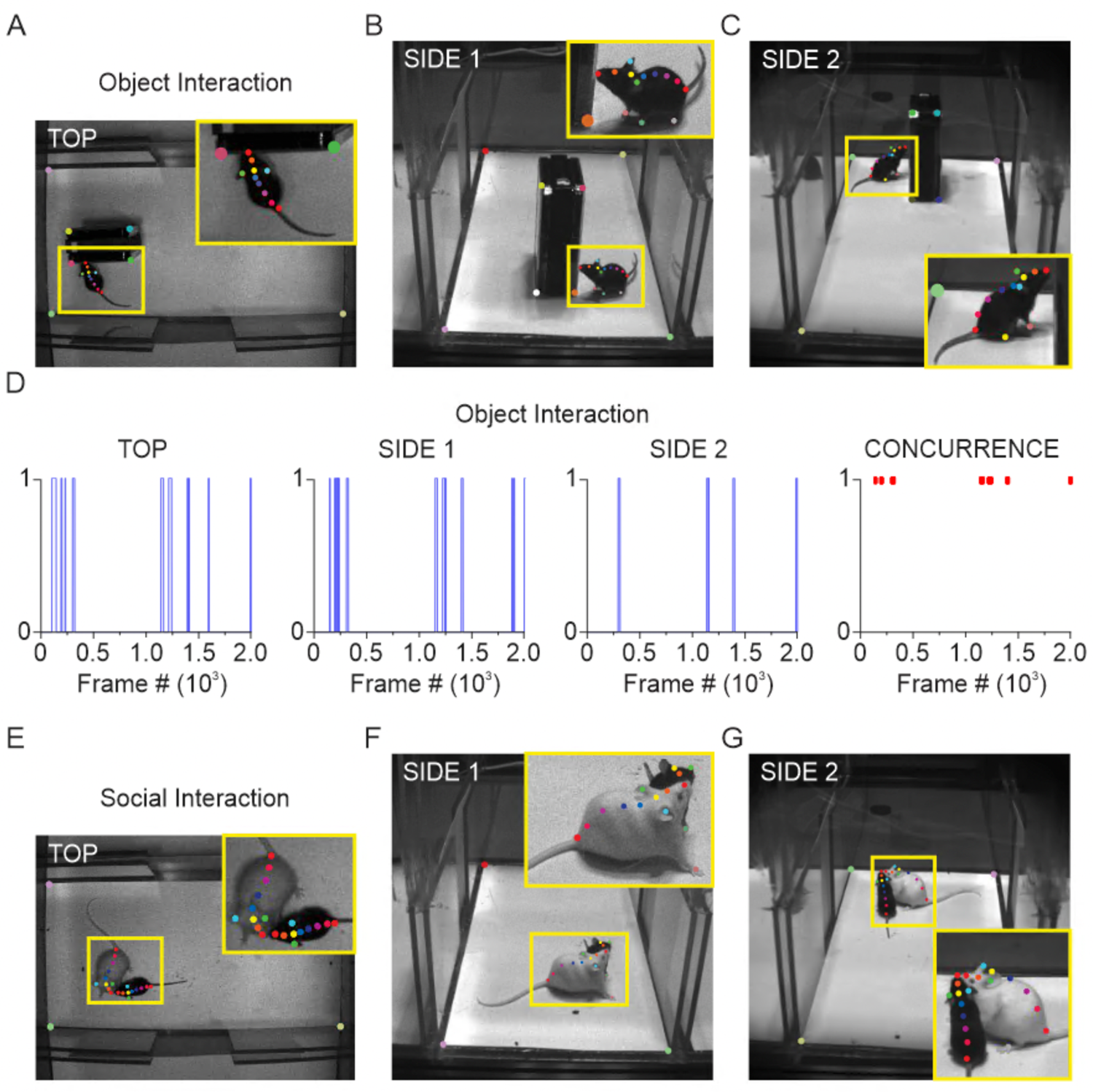
REVEALS can be paired with pose-estimation pipelines to consistently and accurately label mouse body parts and arena features. (**A-C**) Object interaction assay captured from the top (**A**) and two sides (**B, C**); inset zoom in yellow shows body parts labeled by DLC. (**D**) Plots indicating an interaction with the object (binary, 1) within approximately 70 s (2000 frames) recording of the object interaction assay as captured by all three angles. Concurrence (right-most plot) indicates the frames in which at least two cameras captured an interaction. (**E-G**) Social interaction assay recorded from the top (**E**) and two sides (**F**, **G**); inset zoom in yellow shows body parts labeled by DLC. **DLC Annotations. Mouse (snout to tailbase):** red (snout), orange (nose bridge), yellow (head), green (left ear), cyan (right ear), dark blue (neck), violet (body point 1), magenta (centroid), pink (body point 2), red (tailbase). **Object:** yellow (top left corner), cyan (top right corner), pink (bottom left corner), green (bottom right corner), ground top left (white) and ground bottom left (orange) seen in Side 1, ground top right (violet) and ground bottom right (dark green) in Side 2. **Arena:** lavender (top left), red (top right), light green (bottom left), light yellow (bottom right).

## DISCUSSION

We outline the GUI implementation of REVEALS, including its camera control software and video recording functionality. We validate the GUI’s stability, reliability, and accuracy through results showcasing its capability to capture and analyze rodent behavior using DLC pose estimation in an object and social interaction assay. Additionally, the software can be seamlessly integrated into existing DLC and MoSeq pipelines, facilitating the analysis of intricate behaviors.

REVEALS is cost-effective and doesn’t require significant financial resources, which can be particularly important for researchers with budget constraints. REVEALS is accessible, which makes it suitable for many researchers without extensive technical expertise. REVEALS offers tight control over multi-camera frame rates, while keeping them all in sync. In addition, it provides frame timestamps for every camera to allow for post-recording synchronization. Because the images are kept locally rather than through camera storage, dropped frames are a rarity, and as such their scarcity renders the problem negligible in most behavioral experiments.

In future updates, the GUI will have options to change video encoding format as well as adjust output file format, making it possible to reach 120 FPS recordings to allow for capturing of more subtle/rapid movements, such as head shakes (Halberstadt & Geyer, 2013) and discrete paw movement (Forys et al., 2020), without the need for a high-end computer system. The output from REVEALS can be used in popular, easy-to-access software like ImageJ (NIH, Bethesda) without any additional processing, and REVEALS can be operated using a two-step installation protocol for users of any level.

In essence, REVEALS offers a single- or multi-perspective interface for gathering behavioral data, which, when coupled with deep learning algorithms, can enable the scientific community to uncover and characterize complex behavioral traits, ultimately enhancing our comprehension of brain function.

## Supporting information

REVEALS_230822_Supp

## ACKNOWLEDGMENTS

This work was supported by a National Institutes of Health R01 (NIH, 1R01MH129732-01) to A.C-M.; a Brenton R. Lutz award to R.A.P.; and a Boston University Summer Undergraduate Research Fellowship (SURF) to N.M.P-L. We thank Sarah Melzer and members of the Cruz-Martín lab for critical reading of the manuscript and helpful discussions.

## AUTHOR CONTRIBUTIONS

RP: Conceptualization: formulated composition, goals, and scope of the paper and approaches for analyses, Writing—Original Draft: wrote some of parts of original and revised draft. Writing—Review & Editing: editing and feedback for original and revised draft. Visualization: figure design and generation. Wrote and edited original and revised computer code. Experiments: performed behavioral experiments. Analysis: performed DLC analysis.

AW: Writing—Wrote and edited original and revised computer code. Writing—Review & Editing: editing and feedback for original and revised draft.

LF: Conceptualization: formulated composition, goals, and scope of the paper and approaches for analyses, Writing—Original Draft: wrote some of parts of original and revised draft. Writing—Review & Editing: editing and feedback for original and revised draft. Visualization: figure design and generation. Experiments: performed behavioral experiments. Analysis: performed DLC analysis.

NMP-L: Experiments: performed behavioral experiments. Analysis: performed DLC analysis.

MS: Experiments: performed performance analysis. Analysis: performed DLC analysis. Review & Editing: editing and feedback for original and revised draft.

AC-M: Conceptualization: formulated composition, goals, and scope of the paper and approaches for analyses, Writing—Original Draft: wrote some of parts of original and revised draft. Writing—Review & Editing: editing and feedback for original and revised draft. Visualization: figure design and generation. Supervision: mentorship and oversight of the project. Project Administration: management and coordination. Funding acquisition.

## Notes

https://github.com/CruzMartinLab/REVEALS

### Competing Interest Statement

The authors have declared no competing interest.

