## Supplementary material for "REVEALS: An Open Source Multi Camera GUI For Rodent Behavior Acquisition": REVEALS_230822_Supp

A

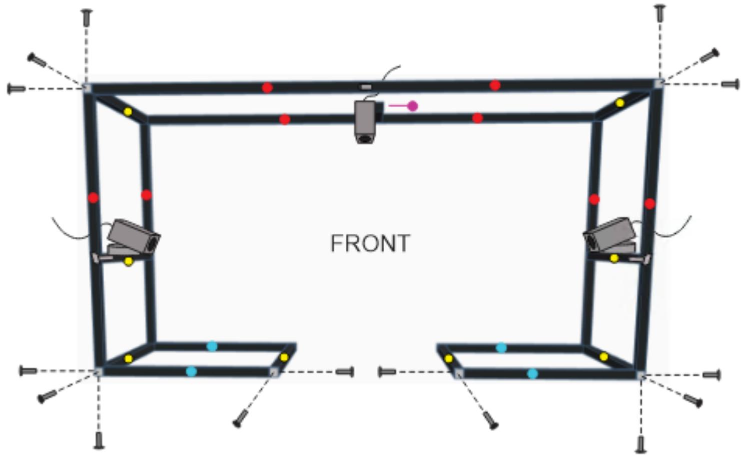


C

| 25mm Square Construction Rail | | | |
|---|---|---|---|
| | 33" | x 8 | XE25L33 |
| | 28" | x 8 | XE25L28 |
| | 21" | x 4 | XE25L21 |
| | 12" | x 1 | XE25L12 |
| Other Components | | | |
| | 25mm | x 12 | XE25W3 |
| | 5/8" | x 32 | SH25LP63 |
| | 1/2" | x 3 | XE25H025 |
| | 1/4" | x 3 | SS25S10 |
| | 1/4" | x 2 | Camera Mount |

B

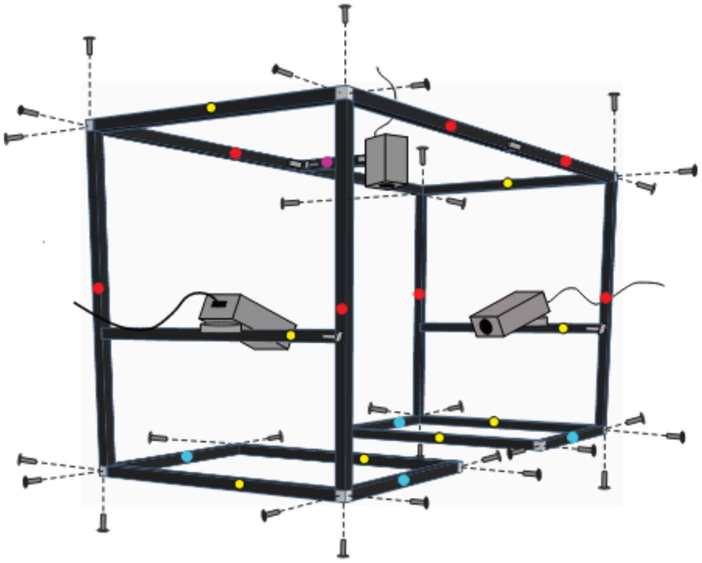

D

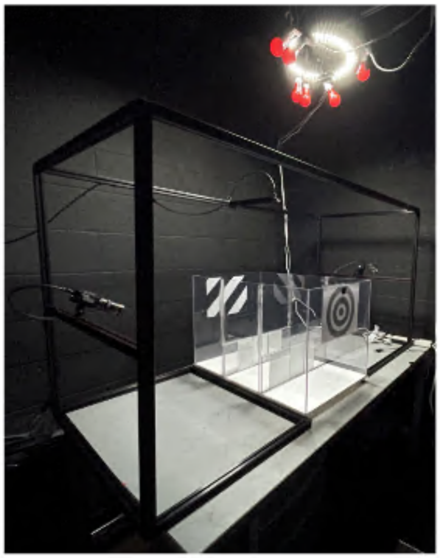

**Supplementary Figure 1. Multi-Camera Recording Rig**. (**A** and **B**) Front (A) and 60° (B) view of recording rig with all parts and accessories needed to build the structure labeled. (**C**) Parts list corresponding to labeled components on schematics of A and B. (**D**) Real picture of recording set up with cameras and behavioral arena in place.

**Supplementary table 1: Key resources table**

| **REAGENT or RESOURCE** | **SOURCE** | **IDENTIFIER** |
|---|---|---|
| **Cameras** | | |
| GS3-U3-32S4M-C 1/1.8" FLIR Grasshopper®3 High Performance USB 3.0 Monochrome Camera | Edmund Optics | 33-534 |
| Flea®3 FL3-U3-32S2M-CS 1/2.8" Monochrome USB 3.0 Camera | Edmund Optics | 86-765 |
| Firefly S FFY-U3-04S2M-S: 0.4 MP, 121 FPS, Sony IMX297, Mono | Teledyne FLIR | FY-U3-04S2M-S |
| **Software and Algorithms** | | |
| REVEALS | https://github.com/CruzMartinLab | N/A |
| Anaconda Software Distribution 2020 | Anaconda Inc. | Anaconda3_64x |
| Spinnaker SDK 2023 | Teledyne FLIR | SpinnakerSDK_FULL_3.0.0.118_x64 |
| **Multi-Camera Recording Structural Rig** | | |
| 25 mm Square Construction Rail, 33" Long, 1/4"-20 Taps | Thorlabs Inc. | XE25L33 |

| | | |
|---|---|---|
| 25 mm Square Construction Rail, 28" Long, 1/4"-20 Taps | Thorlabs Inc. | XE25L28 |
| 25 mm Square Construction Rail, 21" Long, 1/4"-20 Taps | Thorlabs Inc. | XE25L21 |
| 25 mm Square Construction Rail, 12" Long, 1/4"-20 Taps | Thorlabs Inc. | XE25L12 |
| Quick Corner Cube for 25 mm Rails | Thorlabs Inc. | XE25W3 |
| 1/4"-20 Low-Profile Channel Screw, 5/8" Long, 50 Pack | Thorlabs Inc. | SH25LP63 |
| 1/4"-20 Hammerhead Screw, 1/2" Long, 10 Pack | Thorlabs Inc. | XE25H025 |
| 1/4"-20 Set screw, 1” Long, 25 Pack | Thorlabs Inc. | SS25S10 |
| NEEWER 1/4" Mini Cold Shoe Mount Adapter | Neewer, Amazon | B0B11F145V |
